# Examining the Survival of A(H5N1) Influenza Virus in Thermised Whole Cow Milk

**DOI:** 10.1101/2025.03.05.641644

**Authors:** Neda Nasheri, Tamiru Alkie, Todd Cutts, Lemarie Pama, Pablo Romero-Barrios, Angela Catford, Kathleen Hooper-McGrevy, Yohannes Berhane

## Abstract

The recent spillover events of highly pathogenic avian influenza (HPAI) A(H5N1) clade 2.3.4.4b to dairy cattle, and high viral shedding in the milk from infected animals, has created concern that milk and dairy products could be a route for human infection. It has been demonstrated that pasteurization is effective in inactivating A(H5N1) in milk. However, multiple dairy products are made with unpasteurized but thermised milk. The aim of this study was to examine whether some conditions commonly used for thermisation are effective against inactivation of A(H5N1) in whole milk. For this purpose, we artificially inoculated whole raw cow milk with 6.5 log_10_ EID_50_ A(H5N1) and heated for 15 seconds at 60°C, 63°C and 66°C, the viral infectivity was tested using embryonated chicken eggs. We observed over 4 and 5 log_10_ reduction in viral infectivity at 60°C and 63°C, respectively. The viral infectivity was reduced to below the detection limit at 66°C. We also calculated the D-values, the time required to reduce the viral titer by one log_10_, for each treatment and as expected, we observed a decrease in D-values with increasing thermisation temperature. These data demonstrate that thermisation is effective in reducing the viral load and thus they allow for informed risk assessment of A(H5N1) contaminated dairy products made from thermized milk.

## Introduction

In January 2024, a single spillover of highly pathogenic avian influenza HPAI A(H5N1) clade 2.3.4.4b genotype B3.13, from wild birds into dairy cattle in the USA led to multi-state outbreaks, involving over 1000 farms and infecting over 40 humans (1). This incident marked a critical point in understanding the zoonotic potential of A(H5N1) transmission. A second spillover event was reported in January 2025 with A(H5N1) clade 2.3.4.4b genotype D1.1 (2), followed by a third independent transmission with genotype D1.1 in February 2025 (3), demonstrating that transmission of A(H5N1) into dairy cattle can not be considered an improbable event, but rather a potentially recurring threat to the agricultural and public health sectors.

The milk from infected cows with both genotypes can contain high viral titers and thus, concerns have been raised about the safety of milk due to the elevated levels of the virus found in milk from infected cows (4). There is now ample evidence that standard pasteurization heat treatments inactivate the HPAI virus and ensure milk safety (5, 6), so the attention of food safety experts is now turning to the production of dairy products made with unpasteurized milk, including cheeses made with raw or thermised milk. As an initial step in the cheese-making process milk can be heated to temperatures below pasteurization, to reduce the levels of spoilage bacteria, in a process called thermisation. Thermisation temperatures can range from 57°C for 30 minutes to 68°C for a few seconds (7). In this study, we set out to evaluate the inactivation efficiency of three thermisation conditions commonly used in Canada, 66°C, 63°C and 60°C for 15 seconds, against A(H5N1) clade 2.3.4.4.b in whole raw cow milk.

## Materials and Methods

### Virus

The virus used in this study was designated as A/Turkey Vulture/Ontario/FAV473-3/2022 having a stock titer of 1 × 10^8.3^ chicken embryo infectious dose (EID)_50_/mL isolated from Canada and archived at the biosafety level 3+ (BSL-3+) facility at the National Centre for Foreign Animal Disease (NCFAD) in Winnipeg, Manitoba.

### Heat Treatment of the Milk Samples

Fresh bulk raw cow milk was obtained from the University of Manitoba, Department of Animal Science and was treated with an antibiotic/antifungal compound (streptomycin, penicillin, vancomycin, nystatin and amphotericin) (Sigma-Aldrich) for 1 hour at room temperature. The sterility of the milk samples was confirmed by inoculating treated milk onto Columbia Blood Agar Base medium and monitoring daily for microbial growth.

For the heat treatment assays, whole milk was spiked with virus and heated to either 60°C, 63°C, or 66°C for 15 seconds to determine the effects of time and temperature against virus survival. Briefly, three tubes containing 990 μl of antibiotic treated raw non-homogenized milk were placed in a heating block (Eppendorf Thermomixer model R). As described previously (5), a single tube was selected to monitor the milk temperature by inserting an electronic temperature probe, and the temperature adjusted prior to the addition of the viral agent. The remaining two tubes were spiked with 10 μL of viral stock (8.47 log EID_50_ /ml), exposed for 15 seconds, and placed instantly on ice to cool. The virus-spiked, heat treated milk samples were ten-fold serially diluted in fresh antibiotic/antifungal treated raw cow milk and inoculated into 9-days-old embryonated specific-pathogen-free (SPF) chicken eggs (ECEs) (n = 5 eggs/dilution and 200 μL per egg) by following the standard protocol described in the Manual of Diagnostic Tests and Vaccines for Terrestrial Animals (8). The eggs were then incubated at 37°C for 6 to 7 days and candled daily for embryonic mortality as previously described (5). The content of the fourth tube (replicate 4) was ten-fold serially diluted and inoculated into ECEs (n = 5 eggs/dilution) and served as an inoculum titration control to determine the viral load used in each treatment. Four independent experiments were conducted for each temperature (60°C, 63°C or 66°C) for 15 seconds.

### D-value calculation

The D-values (decimal reduction time), which is a measure of the thermal resistance of a microorganism, represents the time required at a specific temperature to reduce the titer by one log unit and is calculated using this formula (7):

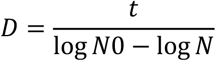

Where t is the time of exposure at a constant temperature, N0 is the initial concentration of the virus and N is the final concentration of the virus after time t.

### Statistical analysis

Statistical analysis was performed using GraphPad Prism v9.0 (GraphPad Software) and the one-way ANOVA method. *P* < 0.05 indicates a significant difference in virus titers in milk samples treated with different thermisation temperatures.

## Results and Discussion

In order to examine the effectiveness of common thermisation conditions on inactivation of A(H5N1), the artificially inoculated whole raw cow milk was treated at 60°C, 63°C and 66°C for 15 seconds and the viral titers after each treatment were determined. As shown in Figure 1, the initial inoculum averaged 6.5 ± 0.56 log_10_ EID_50_/mL, with a reduction to 1.83 ± 1.33 log_10_ and 1.1 ± 0.8 log_10_ EID_50_/mL after exposure to 60°C and 63°C for 15 seconds, respectively. No infectious virus was detectable in inoculated milk samples treated at 66°C for 15 seconds, which translates into approximately 6.5 log_10_ reduction.

**Figure 1.**
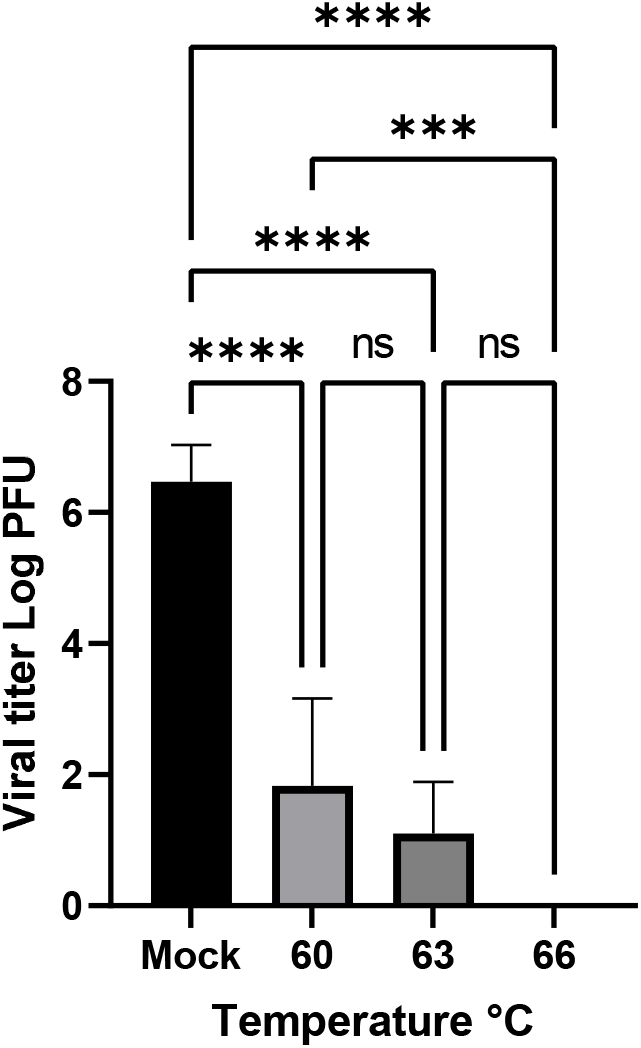
Determination of A(H5N1) viral titers following heat treatment of artificially inoculated raw whole cow milk for 15 seconds at the indicated temperatures. Mock is untreated control. Data shown is the average of 8 replicates from 4 independent experiments. Error bars represent the standard deviation. ns: not significant, ****P < 0.0001, ***P < 0.001 calculated by one-way ANOVA.

Statistically, the reductions in viral titers at thermisation temperatures (60°C, 63°C, and 66°C) were significantly different from the untreated control, while the reductions in viral infectivity between 60°C and 63°C, and 63°C and 66°C, were not significantly different from each other.

Next, we calculated the D-values to determine the time that is required to achieve 1 log reduction at each temperature. As expected, the D-values decreased with increasing temperature, indicating a more rapid rate of viral inactivation at higher thermisation temperatures. The D-values calculated for 60°C, 63°C, and 66°C were 3.24 seconds, 2.82 seconds, and 2.34 seconds, respectively (Table 1). Overall, these results demonstrate that thermisation of whole cow milk contaminated with A(H5N1) virus at temperatures above 60°C could considerably reduce the viral load.

**Table 1.** D-values calculated for A (H5N1) in raw whole cow milk after treatments for 15 seconds.

| Temperature | D-values |
| --- | --- |
| 60°C | 3.24 s |
| 63°C | 2.82 s |
| 66°C | 2.34 s |

One limitation of this study is that the heating treatments were conducted under laboratory settings and not using the equipment and milk volumes used by the dairy industry. Furthermore, the A(H5N1) transmissibility and the potential infectious dose of A(H5N1) to humans through ingestion is unknown, however, studies on cynomolgus macaques revealed that inoculation via ingestion can cause infection, albeit with milder symptoms compared to inoculation via respiratory routes (9). On the other hand, drinking A(H5N1) contaminated raw milk has been demonstrated to lead to severe infection in mice and cats (10, 11) indicating that different species may exhibit varying degrees of susceptibility to A(H5N1)depending on the exposure route. Furthermore, viral inactivation efficacy may vary by strain, as some A(H5N1) genotypes may demonstrate more resistance to heat treatment than others. Notably, A(H5N1) clade 2.3.4.4b virus has been observed to have enhanced environmental stability, which may complicate pathogen control in dairy products derived from infected cattle (12).

This study underscores the relevance of thermisation as a potential mitigation strategy for A(H5N1) in the dairy industry. In the absence of data about the effect of the different stages of cheese processing on the survival of the HPAI virus, cheese made with thermised milk can be considered less likely to lead to exposure to the HPAI virus compared with raw milk cheese.

Given the increasing frequency of A(H5N1) spillover events and the documented potential for zoonotic transmission via contaminated dairy products, these findings underscore the need for a more comprehensive risk assessment regarding A(H5N1) contamination in dairy products. Future studies should aim to evaluate A(H5N1) survival through various stages of dairy processing, including during fermentation, aging, and storage.

## Acknowledgements

This study is financially supported by the Defence Research and Development Canada (DRDC) CW2248375 -39903-23076 CSSP 2541 and CFIA HPAI Emergency Grant as well as the Bureau of Microbial Hazards at Health Canada.

## Conflict of Interest

The authors declare no conflict of interest.

## Notes

### Competing Interest Statement

The authors have declared no competing interest.

